# Cell-intrinsic genetic regulation of peripheral memory-phenotype T cell frequencies

**DOI:** 10.1101/355313

**Authors:** Amanpreet Singh Chawla, Parna Kanodia, Ankur Mukherjee, Vaibhav Jain, Gurvinder Kaur, Poonam Coshic, Kabita Chatterjee, Nitya Wadhwa, Uma Chandra Mouli Natchu, Shailaja Sopory, Shinjini Bhatnagar, Partha P. Majumder, Anna George, Vineeta Bal, Satyajit Rath, Savit B. Prabhu

**Author notes:** These authors contributed equally to the manuscript (Amanpreet Singh Chawla, Parna Kanodia). These authors contributed equally to the manuscript (Anna George, Vineeta Bal, Satyajit Rath, Savit B. Prabhu). **Address for correspondence**: Savit B. Prabhu, Translational Health Science and Technology Institute, NCR Biotech Science Cluster, 3rd Milestone, Faridabad – Gurgaon Expressway, PO box #04, Faridabad – 121001, Haryana, India.

## Abstract

Memory T and B lymphocyte numbers are thought to be regulated by recent and cumulative microbial exposures. We report here that memory-phenotype lymphocyte frequencies in B, CD4 and CD8 T-cells in 3-monthly serial bleeds from healthy young adult humans were relatively stable over a 1-year period, while recently activated -B and -CD4 T cell frequencies were not, suggesting that recent environmental exposures affected steady state levels of recently activated but not of memory lymphocyte subsets. Frequencies of memory B and CD4 T cells were not correlated, suggesting that variation in them was unlikely to be determined by cumulative antigenic exposures. Immunophenotyping of adult siblings showed high concordance in memory, but not of recently activated lymphocyte subsets, suggesting genetic regulation of memory lymphocyte frequencies. To explore this possibility further, we screened effector memory (EM)-phenotype T cell frequencies in common independent inbred mice strains. Using two pairs from these strains that differed predominantly in either CD4EM and/or CD8EM frequencies, we constructed bi-parental bone marrow chimeras in F1 recipient mice, and found that memory T cell frequencies in recipient mice were determined by donor genotypes. Together, these data suggest cell-autonomous determination of memory T niche size, and suggest mechanisms maintaining immune variability.

## Introduction

Gene-environment interplay in immune phenotypes has been extensively studied using steady-state cellular immune profiles (1–3), functional immune responses (2,4–8), post-vaccination responses (9,10) and V,D and J usage biases in naïve and memory T and B cell compartments (11). Some of these studies have identified genomic correlates associated with specific steady-state immune phenotypes (2) and vaccine responses (12). However, there are conflicting findings regarding the relative importance of genetic versus environmental factors in regulation of immune phenotype (1,2,13,14), warranting further investigations at the population-level in humans and mechanistic studies in mice on regulation of individual immune phenotypes.

Memory subsets in T and B lymphocytes are immune populations that are generated in response to past immunogen exposure. Immunological memory, providing long-term persistence of antigen-experienced cells contributing to rapid and robust responses following re-exposure (15), is likely to have evolved in an ecosystem where environmental challenges including repeated infections would be the norm (16) and the persistence of long-lasting antigen-specific cells generated during immune responses would confer survival advantage (17). However, it is possible that larger memory lymphocyte pool sizes may carry costs such as restriction of space for the more repertoire-diverse naïve T cell compartment (18), attrition of pre-existing memory (19), and other bioenergetic costs (20). Such selection may well result in ‘optimum’ sizes of memory lymphocyte pools (21), and these pool sizes could show population diversity, depending on the diversity of pathogens in the ecosystem and their exposure rates (22). Diversity in pool sizes of memory lymphocytes in a population could thus be determined by a combination of genetic variability and diversity of environmental exposures.

A number of mechanisms can be envisaged regulating the pool size of the memory T cell compartment, including cumulative life-time antigen-exposure and re-exposure, antigenic persistence, degree of expansion, cell survival, attrition and niche-space availability (21,23–26). Immune cells occupy a limited niche space in lymphoid organs (27) or in the periphery (28). This niche size could be a function of size and/or structure of supporting lymphoid tissue architecture (29), along with intrinsic properties of cells occupying the niche. Similar determinants could affect steady state levels of transient cell populations such as immediate-effector T cells (30) and plasmablasts, but their steady state levels would be expected to fluctuate more with short-term environmental changes.

On this background, we have explored the possibility of gene-environment interplay affecting the steady state pool sizes of lymphocytes post-activation. We have used peripheral blood leucocyte immunophenotyping in serial bleeds from healthy young adult human volunteers to assess inter-individual and temporal variability in lymphocyte memory and effector compartments, and report that while both categories showed inter-individual variability, temporal variability was far less in memory than in effector subsets. We have next used peripheral blood leucocyte immunophenotyping of human siblings, and report that memory lymphocyte subset pool sizes were far closer between siblings than between non-siblings. We then further confirmed a role for genetic factors in determining memory lymphocyte pool sizes in independent inbred mouse strains, in which bi-parental mixed bone marrow chimeras demonstrated a parental genotype-driven inheritance of CD4 and CD8 T cell memory pool sizes. Together, our data from both humans and mice provide novel insights in the determination of memory lymphocyte pool size.

## Materials and methods

### Ethics approval and consent to participate

#### Human

Informed written consent was obtained from all human volunteers who participated in the study. The study was approved by ethics committees of National Institute of Immunology (approval number IHEC/AKS/45/2013) and All India Institute of Medical Sciences (approval number IEC/NP-471/2013). The methodologies used in the study were in accordance with approved guidelines. All experimental protocols used in this study were approved by Institutional Ethics Committee (Human Research) of National Institute of Immunology and All India Institute of Medical Sciences.

#### Animal

Mice were maintained and used according to the relevant rules and regulations of Government of India and with the approval of Institutional Animal Ethics Committee, National Institute of Immunology (approval number 381/15). All mice experimental protocols were in accordance with approved guidelines and were approved by Institutional Animal Ethics Committee of National Institute of Immunology.

### Human subjects and blood collection

Forty-five healthy adult volunteers were recruited into the study and were bled 3-monthly for 12 months (total of 4 bleeds). Individuals with acute or chronic illness, medication, recent vaccination or pregnancy in women were excluded. Ten ml heparinized peripheral blood was collected from healthy subjects after informed written consent. The study was approved by institutional ethics committee of National Institute of Immunology (NII). All protocols were in accordance with approved guidelines. For sibling study, 37 families were recruited, in which two or more siblings were willing for participation. The family-based study was approved by institutional ethics committees of NII and All India Institute of Medical Sciences (AIIMS).

Exclusion criteria were similar to that mentioned above. For looking at correlations between the various memory and effector cell subsets, we utilized the published raw data from a larger (n=71) cohort of adult healthy volunteers who participated in a previous study (31).

### Sample processing and flow cytometry of human samples

Blood samples were processed without delay (less than three hours). Total leukocyte counts and differential counts were obtained from standard hematologic methods. Peripheral blood mononuclear cells (PBMCs) were separated by density gradient centrifugation. PBMCs were washed, counted and divided into aliquots of about 1 million cells per ml per vial and cryopreserved in 10% DMSO in bovine serum until assays were performed. For flow cytometry, PBMCs were thawed, washed and incubated with the following antibodies organized into two cocktails: The T cell cocktails consisted of CD3 (UCHT1, eBioscience), CD4 (OKT4, eBioscience), CD8 (SK1, eBioscience), CD45RO (UCHL1, eBioscience), CCR7 (150503, BD) and the B cell cocktail consisted of CD19 (SJ25C1, BD), CD20 (2H7, eBioscience), CD38 (HIT2, eBioscience), CD43 (eBio84-3C1, eBioscience), CD27 (M-T271, eBioscience). A single control sample was run along with all the samples to ensure that gating is comparable across experiments. Samples were acquired in BD Verse and analysis was done using flowjo (Treestar). CD4 and CD8 memory cells were defined as the CD45RO+ fractions (32) of CD8 and CD4 cell compartments. Transient T effector-memory, RA+ve (TEMRA) cells were defined as CD45RO-CCR7-subsets (32) of CD8 cells and of CD4 cells. Memory B cells were defined as CD27+CD43-fraction of the CD19+CD20+ B cell subset (32). Plasmablasts were defined as the CD38+CD20-subset of CD19+ B cells (32).

### Mice

The following strains of mice were used for the study: C57Bl/6J, BALB/cJ, SJL/J, CBA/CaJ, B6.SJL-Ptprca Pepcb/BoyJ (B6.SJL), CB10-H2b/LilMcdJ (BALB/b), FVB/NJ, C3H/HeOuJ. Mice strains used in the study were obtained as breeder stock from the Jackson Laboratory (Bar Harbor, ME) and bred and maintained under specific-pathogen free conditions in the Small Animal Facility of the National Institute of Immunology. All mice were maintained and used according to the relevant rules and regulations of Government of India and with the approval of Institutional Animal Ethics Committee, National Institute of Immunology. All experiments used age and gender matched mice. Littermates were used as controls. Phenotyping of mouse spleen T cells was done using the following antibodies (all from eBioscience): CD4 (RM4-5), CD8 (53-6.7), CD44 (IM7), CD62L (MEL-14), CD25 (PC61.5), CD45.1 (A20) and CD45.2 (104). NK cells were phenotyped using the following antibodies (all from eBioscience): CD90 (53-2.1), B220 (RA3-6B2) and Ly49b (DX-5). Surface staining was done by incubating 1×10^6^ cells in staining buffer (PBS containing 2% BSA and 0.05% NaN3) for 30 mins on ice. The cells were washed thrice with cold staining buffer and samples were acquired on BD Verse and analysis was done using flowjo (Treestar).

### Ex vivo cell preparations

Spleen was dissected from mice euthanized by cervical dislocation and teased between a pair of frosted slides to obtain single cell suspension. Red blood cells in the suspension were lysed by osmotic shock using water, washed and then re-suspended in complete medium.

### Mix bone marrow chimeras

F1 hybrids were irradiated at 800rads in gamma chamber (BARC, Mumbai) with Co^60^ as a source for gamma rays. Bone marrow cells from each parental strain were mixed in 1:2, 1:1 and 2:1 ratios and a total of 15 million cells transferred into irradiated F1 generation mice (n=24 for B6.SJL - CBA/CaJ chimera and n=19 for SJL/J - BALB/cJ chimera) and reconstitution allowed for two months. Based on the ratio of CD45.1 to CD45.2 in reconstituted chimera, biological outliers showing extreme ratios more than 10-fold from the expected were removed from further analysis (4 outliers in B6.SJL - CBA/CaJ chimera and 2 outliers in SJL/J - BALB/cJ chimera were removed). Phenotyping was done by using above mentioned CD markers. Doublets were excluded from the population using height and area parameters of forward and side scatter.

CD4 effector memory cells were defined as the CD4+ CD25-CD44hi CD62Llo fractions and CD8 effector memory cells were defined as the CD8+ CD44hi CD62Llo fractions.

### Statistical analysis

For frequency and cell count comparisons, a non-parametric test (Mann Whitney) was used. Correlations were estimated using Spearman’s correlation coefficient. The Bonferroni method was used to correct for multiple comparisons in testing the significance of elements of the pairwise correlation coefficient matrix. Bootstrapping (resampling) methods were used for comparison of intra-individual variances with between-individual variances and for comparison of sibling pairs vs unrelated pairs. Comparison of multiple inbred mouse strains was done using Analysis of Variance (ANOVA) with post-hoc testing for pair-wise comparison.

### Gene expression analysis

Gene expression (microarray) data of sorted splenic ‘naïve’ CD4+ve CD62L-ve CD4 T cells of mice published previously (33) was obtained from GEO database (accession: GSE60337) and analysed using GEO2R tool. Genes with p-value of less than 0.05 after multiple testing (Benjamini) was considered significantly differentially expressed. Gene Ontology enrichment analysis was done using web-based Gene Ontology enrichment analysis and visualization tool (34) (GORILLA) and Database for Annotation, Visualization and Integrated Discovery (DAVID)(35). To reduce false positive results in enrichment, in all enrichment analyses, the enrichment of candidate genes that were differentially expressed were analysed against a background of all the genes that were used in array experiment.

## Results

### Human peripheral T and B memory cell levels show temporal stability

Firstly, we examined if variation in the memory T and B cell subsets in the population were relatively stable over time, or were significantly affected by fluctuations possibly contributed by short-term environmental exposures. To test this, we characterized the peripheral blood leucocyte subset phenotype of a group of human volunteers [n=45, male: female ratio = 26:19, mean age = 27.16 years (SD = 6.75)] every 3 months for a period of 1 year. Gating strategy for the memory subsets is indicated in supplementary figure 1 (Fig S1).

Representative data of frequencies of each subset over 4 time points are shown in Fig S2 and are shown quantified in Figure 1. Variations of memory B, memory CD4 and memory CD8 frequencies were significantly lower within individuals than between individuals (Fig. 1A, 1B, 1C). On the other hand, intra-individual variances in plasmablasts (Fig. 1D) and CD4 TEMRA cells (Fig. 1E) were no different from inter-individual variances. When absolute cell numbers per μl blood were calculated, broadly similar trends were seen (Fig. 1G, 1H, 1I, 1J, 1K). Curiously, CD8 TEMRA frequencies (Fig. 1F) and counts (Fig. 1L) tended to be similar to CD8 memory (Fig. 1C, 1I) in that they showed lower intra-individual than inter-individual variation. These data suggested that intra-individual short-term fluctuations in memory B and T subsets are unlikely to be a major explanation for the variation that is seen in memory lymphocyte subsets in humans.

**Figure 1:**
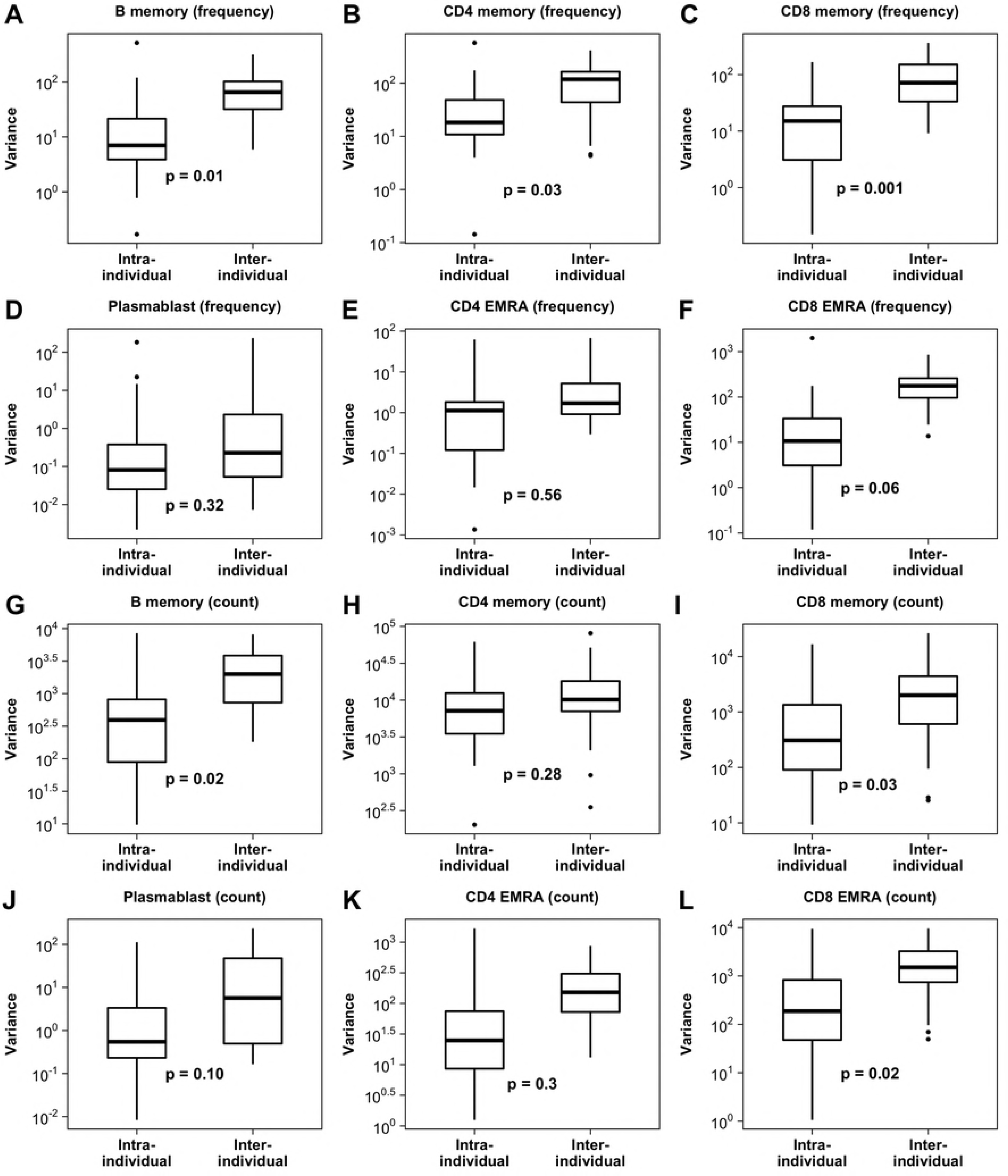
Comparison of intra-individual and inter-individual variance for immune subsets in Humans.

Box plots indicate median and interquartile ranges of variances of cell frequencies and counts in Human volunteers. Upper whisker extends till the highest value that is within 1.5 times the interquartile range from 3rd quartile. Lower whisker extends till the lowest value that is within 1. 5 times the interquartile range from 1st quartile. Outliers are shown as dots. Intra-individual variances indicate variance of subset frequency or count over 4 time points in each individual (n=45). Inter-individual variances indicate variance of subset frequency or count in randomly chosen set of different individuals (n=45). P-values obtained by bootstrapping are as indicated in the panels.

### Correlations between CD4, CD8 and B cell memory

Since memory cells can be long-lived and could have been generated by antigenic exposure in the relatively remote past, it remained possible that cumulative antigen exposures contribute to determining steady-state memory T and B cell levels. Microbial exposure commonly leads to generation of CD4 T-dependent B cell responses, since CD4 T cells are necessary for B cell germinal centre responses and hence vital to memory B cell generation. Thus, it would be expected that there would be coordinated accumulation of memory cells in the CD4 T and B cell compartments leading to a positive correlation between CD4 T memory and B memory cell frequencies, if cumulative antigenic exposures were to contribute substantially to determining memory T and B cell frequencies. However, there was no correlation between CD4 T cell memory frequencies and B cell memory frequencies in a given individual (Fig. 2A). Thus, cumulative antigen exposure may not be a major determinant of memory CD4 and B cell levels. Interestingly, CD4 and CD8 memory T cell subsets showed a strong correlation (correlation coefficient = 0.61, p < 0.001) (Fig. 2B), suggesting shared determinants of memory cell subsets within the T cell lineage. On the other hand, frequencies of CD4 TEMRA or CD8 TEMRA cells or plasmablasts did not show any correlations with memory T or B cell frequencies or with each other (Table 1).

**Table 1:** Correlation coefficient (Spearman) of correlation between cell subsets in adult volunteers.

| Pairs of variables | Correlation coefficient<br>(Spearman) | Adjusted p-value<br>(Bonferroni) |
| --- | --- | --- |
| Plasmablast – B memory | 0.17 | 1 |
| CD4 memory - B memory | 0.0009 | 1 |
| CD4 TEMRA - B memory | -0.006 | 1 |
| CD8 memory - B memory | -0.09 | 1 |
| CD8 TEMRA - B memory | 0.038 | 1 |
| CD4 memory - Plasmablast | -0.04 | 1 |
| CD4 TEMRA - Plasmablast | 0.09 | 1 |
| CD8 memory – Plasmablast | -0.0017 | 1 |
| CD8 TEMRA - Plasmablast | 0.16 | 1 |
| CD4 TEMRA - CD4 memory | -0.26 | 0.335 |
| CD8 memory - CD4 memory | 0.6 | 8.90E-08 |
| CD8 TEMRA - CD4 memory | -0.064 | 1 |
| CD8 memory - CD4 TEMRA | -0.16 | 1 |
| CD8 TEMRA - CD4 TEMRA | 0.32 | 0.065 |
| CD8 TEMRA - CD8 memory | 0.2 | 1 |

**Figure 2:**
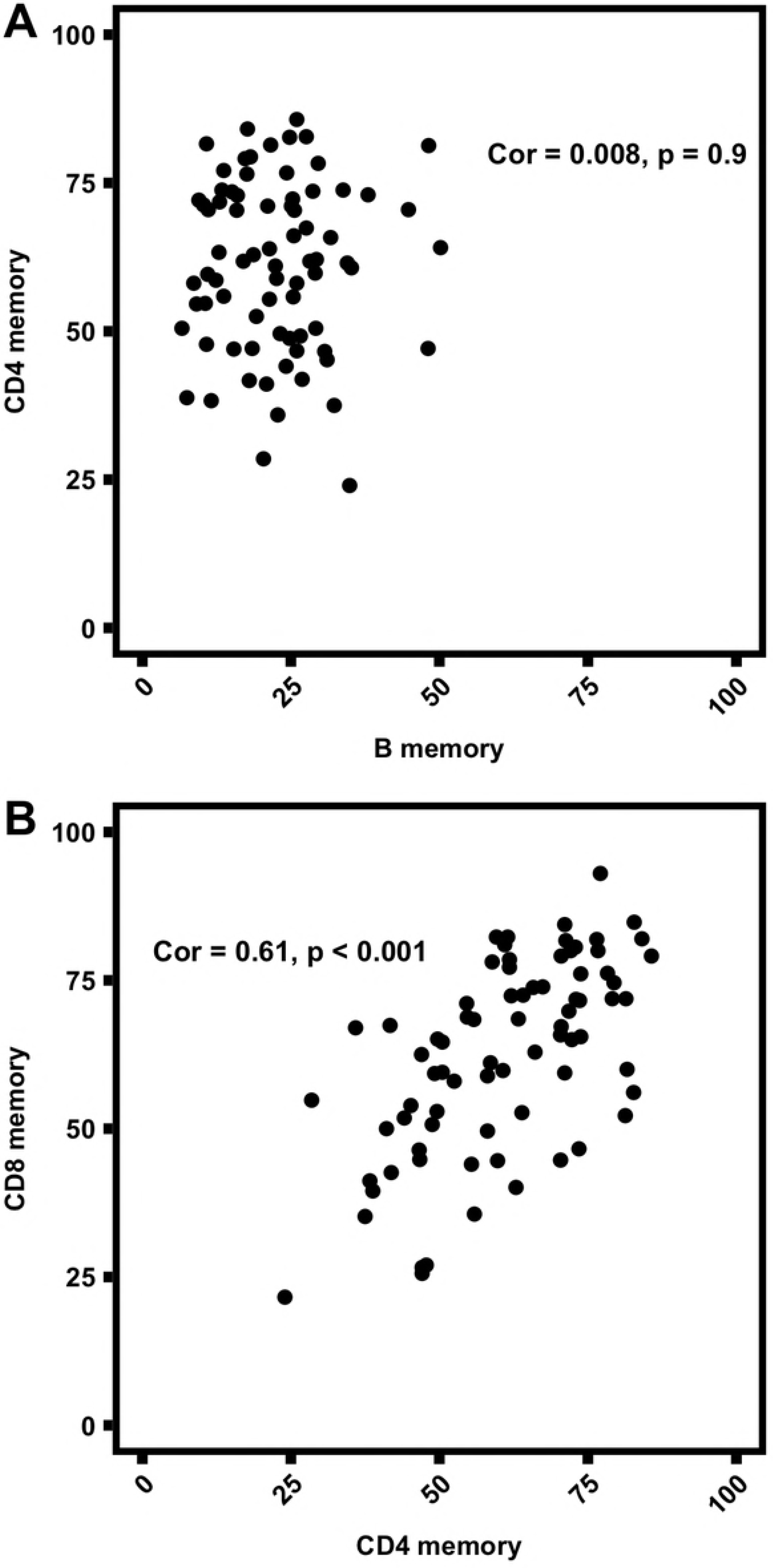
Correlation between memory B and memory CD4 T cell frequencies (A) and between memory CD4 and memory CD8 T cell frequencies (B) Memory CD4, memory CD8 and memory B cell frequencies are expressed as percentage of total CD4, CD8 and B cells respectively. Each dot represents data from one individual donor (n=71). Correlation coefficient (Spearman) and p-value are as indicated.

### Siblings show concordance in T and B cell memory levels

Since these descriptive and correlative data from humans showed associations that suggested that memory phenotypes might be genetically regulated, we explored this further by a family based study where peripheral blood leucocytes from 80 full siblings from 37 families were immunophenotyped [male: female ratio = 38:42, mean age = 31.8 years (SD=12.7)]. We compared the differences in cell frequencies between siblings with the differences between random pairs of non-siblings for memory CD4, memory CD8 and memory B cells, and for plasmablasts, CD4 TEMRA or CD8 TEMRA cell frequencies. The analysis showed that differences in cell frequencies between siblings were significantly less than the differences between random pairs of non-siblings for memory CD4, memory CD8 and memory B cells, but not for plasmablasts, CD4 TEMRA or CD8 TEMRA cell frequencies (Table 2), suggesting that there was indeed likely to be a genetic contribution to the variations in memory cell frequencies, although possible contributions from early life co-habitation could not of course be ruled out.

**Table 2:**
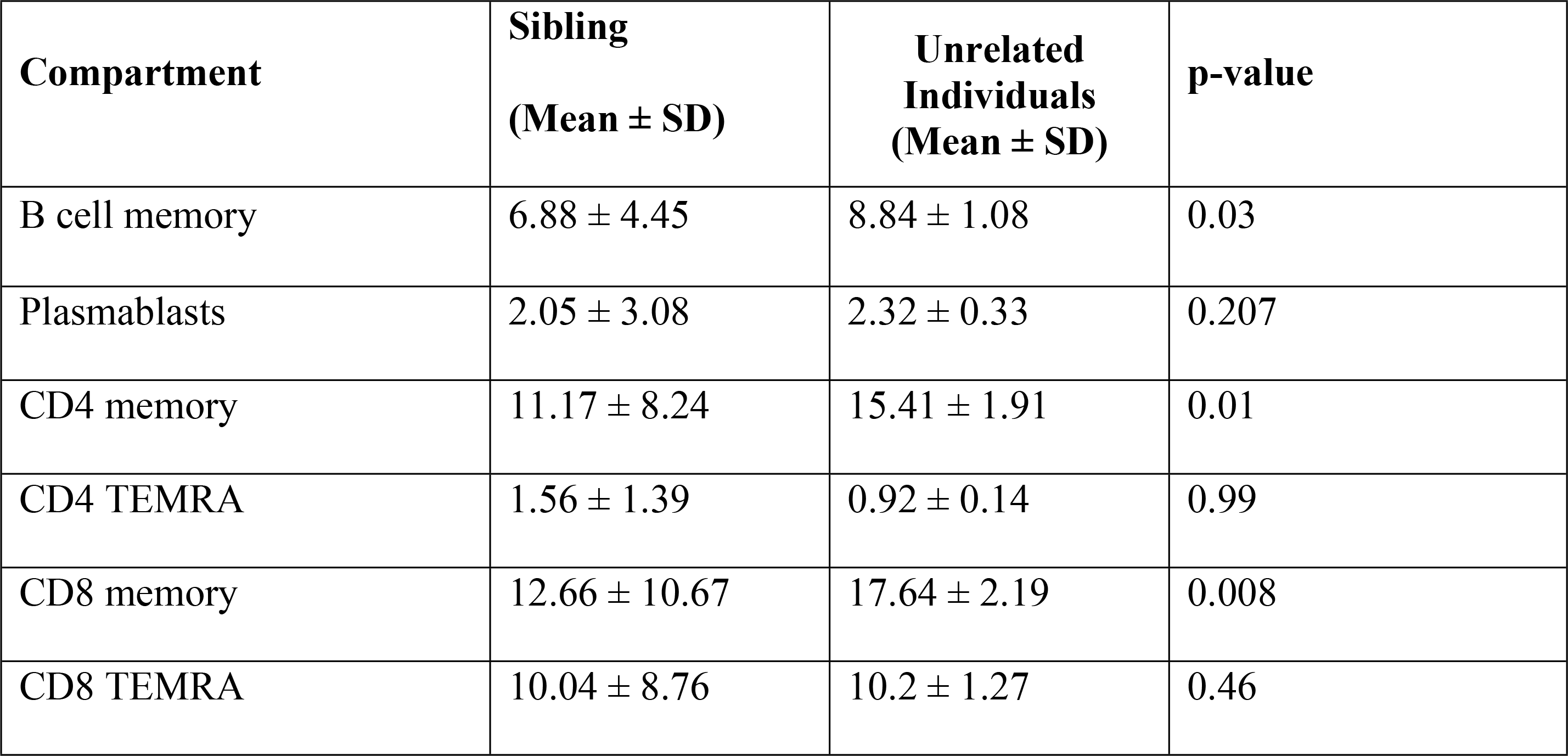
P-values of difference between within-sibling pairs and between-sibling pairs.

| **Compartment** | **Sibling**<br>**(Mean ± SD)** | **Unrelated**<br>**Individuals**<br>**(Mean ± SD)** | **p-value** |
| --- | --- | --- | --- |
| B cell memory | 6.88 ± 4.45 | 8.84 ± 1.08 | 0.03 |
| Plasmablasts | 2.05 ± 3.08 | 2.32 ± 0.33 | 0.207 |
| CD4 memory | 11.17 ± 8.24 | 15.41 ± 1.91 | 0.01 |
| CD4 TEMRA | 1.56 ± 1.39 | 0.92 ± 0.14 | 0.99 |
| CD8 memory | 12.66 ± 10.67 | 17.64 ± 2.19 | 0.008 |
| CD8 TEMRA | 10.04 ± 8.76 | 10.2 ± 1.27 | 0.46 |

### Mouse strains differ in their memory T cell frequencies and show cell-autonomous regulation of memory T cell phenotype

These data from humans indicated the possibility of a genetic contribution to the determination of memory T and B cell pool sizes. To examine this possibility more rigorously, we examined independent inbred strains of mice. Since specific markers for B cell memory are uncertain (36) and since this compartment in mice may be smaller than in humans (37), we restricted these studies to T cell memory subsets alone.

The gating strategy used for defining effector memory (EM) CD4 (Fig 3A to 3D) and CD8 T cells (Fig 3E to 3H) in a group of representative mice available in our laboratory are shown in Figure 3. EM populations in CD4 and CD8 subsets were distinguished by unambiguous contours, whereas in some strains (eg. BALB/cJ) the central memory (CM) populations were difficult to gate out from the CD44 negative naïve pool using the conventional markers of memory (CD44 and CD62L). CM phenotype CD4 T cells in the BALB/c strain have been previously shown to contain recent thymic emigrants with a CD44-high phenotype (38). CM phenotype CD8 T cells have also been reported to contain antigen-inexperienced, non-memory (“virtual memory”) cell types (39) as well as regulatory CD8 T cells (40), further confounding our analysis of memory subsets. Hence, we have avoided CM phenotype T cells and focused on the unambiguously defined EM populations for interpreting differences in memory between the genetically different strains.

**Figure 3:**
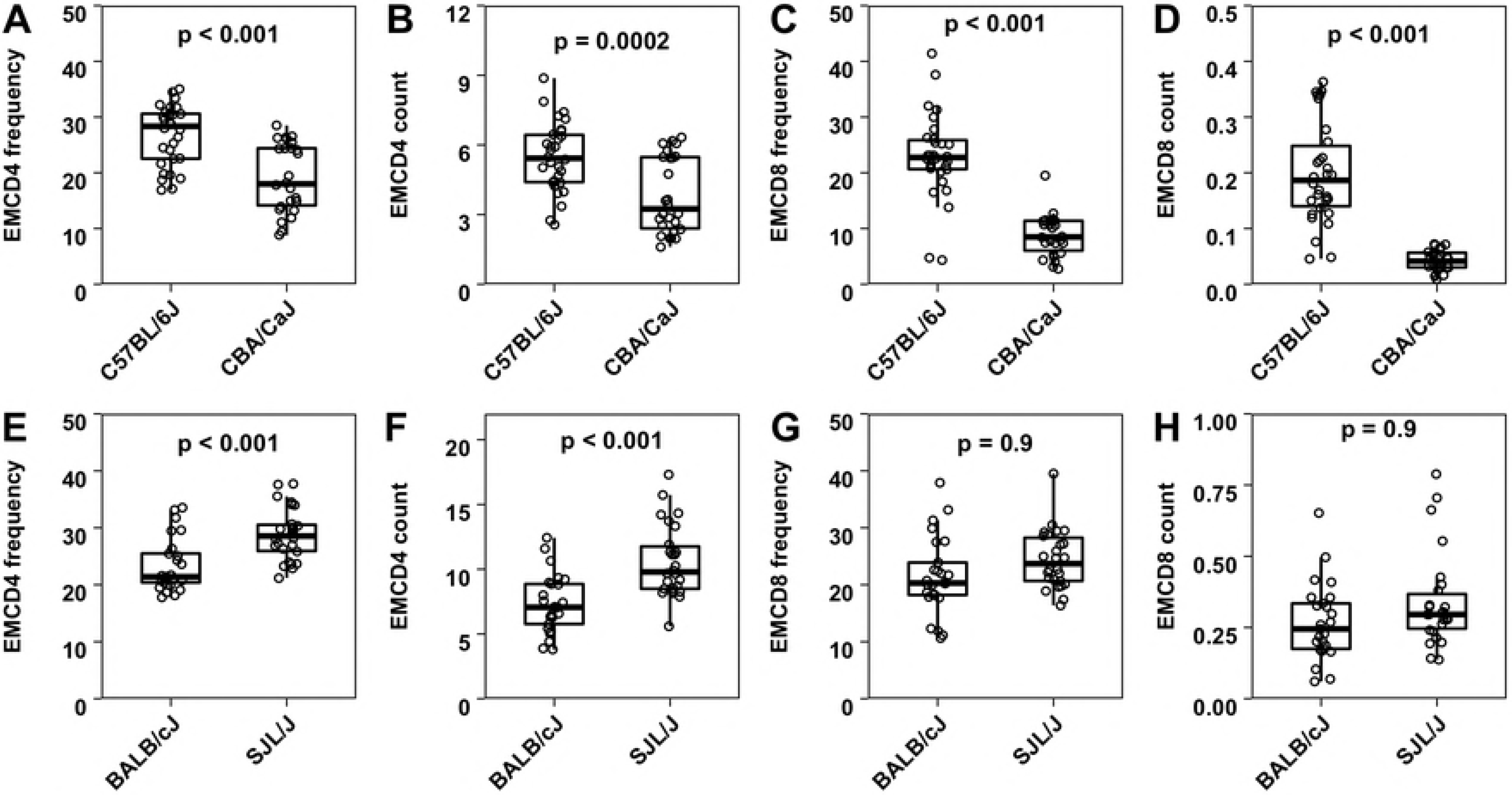
Gating strategy for memory subsets in mouse splenic CD4 T cells (A to D) and CD8 T cells (E to H) Each plot shows representative gates from a single mouse strain as indicated. CD4 memory subsets are gated on conventional CD4+ CD25-gate. CD8 memory subsets are gated on CD8+ gate.

Our preliminary data showed that CBA/CaJ and C57BL/6J showed differences in both EMCD4 and EMCD8 compartments with C57BL/6J showing higher EMCD4 and EMCD8 frequencies and counts than CBA/CaJ (Fig 4A - 4D). On the other hand, BALB/cJ and SJL/J differed in EMCD4 but not EMCD8 compartment (Fig 4E - 4H), with SJL/J showing significantly higher EMCD4 frequencies and counts than BALB/cJ (Fig 4E, 4F).

**Figure 4:**
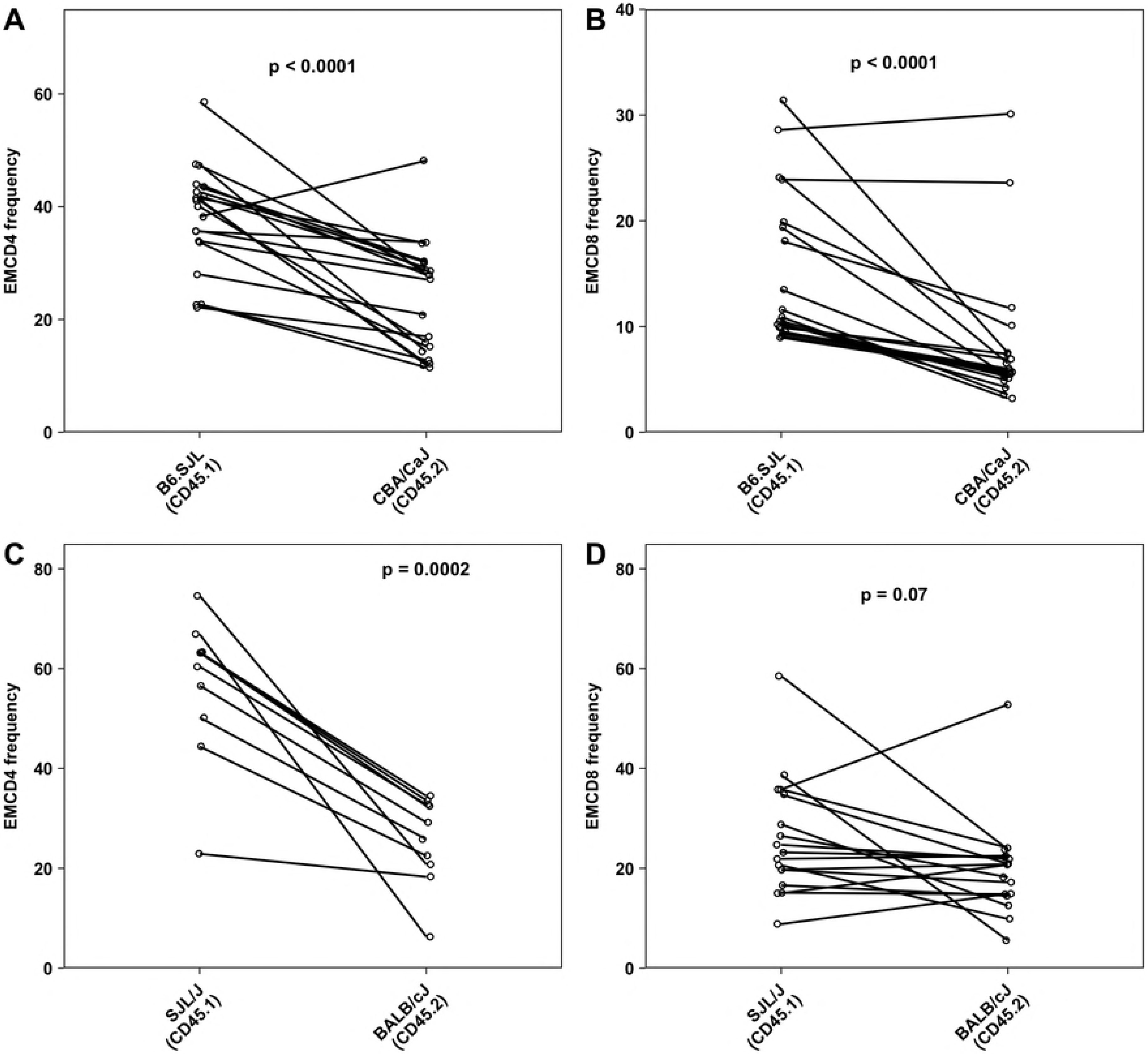
EMCD4 and EMCD8 frequencies and counts (in millions) in spleen in C57BL/6J versus CBA/CaJ (A to D) and BALB/cJ versus SJL/J (E to H) Each dot represents data from one individual mouse (n > 25 per group). EMCD4 and EMCD8 frequencies indicate frequencies of EM compartment out of total conventional CD4 (CD4+CD25−) and CD8 T cells respectively. Cell counts are expressed in millions. Box plots indicate median and interquartile ranges. Upper whisker extends till the highest value that is within 1.5 times the interquartile range from 3rd quartile. Lower whisker extends till the lowest value that is within 1.5 times the interquartile range from 1st quartile. The p-values are as indicated in the panels.

To test the source of the differences observed in EMCD4 frequencies between the strains, we prepared bi-parental mixed bone marrow chimeras in which bone marrow cells from both parents, in various ratios, were transferred into irradiated F1 mice. We used heavy irradiation that is reported to successfully engraft parental donor cells in F1 recipients (41–44) and ensured endogenous NK cell depletion post-irradiation (Fig S3 A and B). We also confirmed that chimerism is successful by comparing injected donor cell ratios with ratios of CD45.1 to CD45.2 lymphocytes in reconstituted F1 mice (Fig S3 C and D) and found no statistically significant differences, ruling out the possibility of unequal reconstitution by donor strains. The mean frequency of endogenous cells was 10.1 % in B6.SJL - CBA/CaJ chimera (SD-6.5) and 25.9 % in BALB/cJ - SJL/J chimera (SD-12.3). After waiting two months for generation of memory cells from the donor genotypes, these mice were phenotyped for CD4 and CD8 EM compartments. Memory CD4 and CD8 frequencies of the donor-derived cell population (Fig 5A to 5D) in these chimeras showed similar trends as that in parental strains (Fig 4). These results suggest that genetic factors contribute in cell intrinsic fashion to determining EM T cell frequencies.

**Figure 5:**
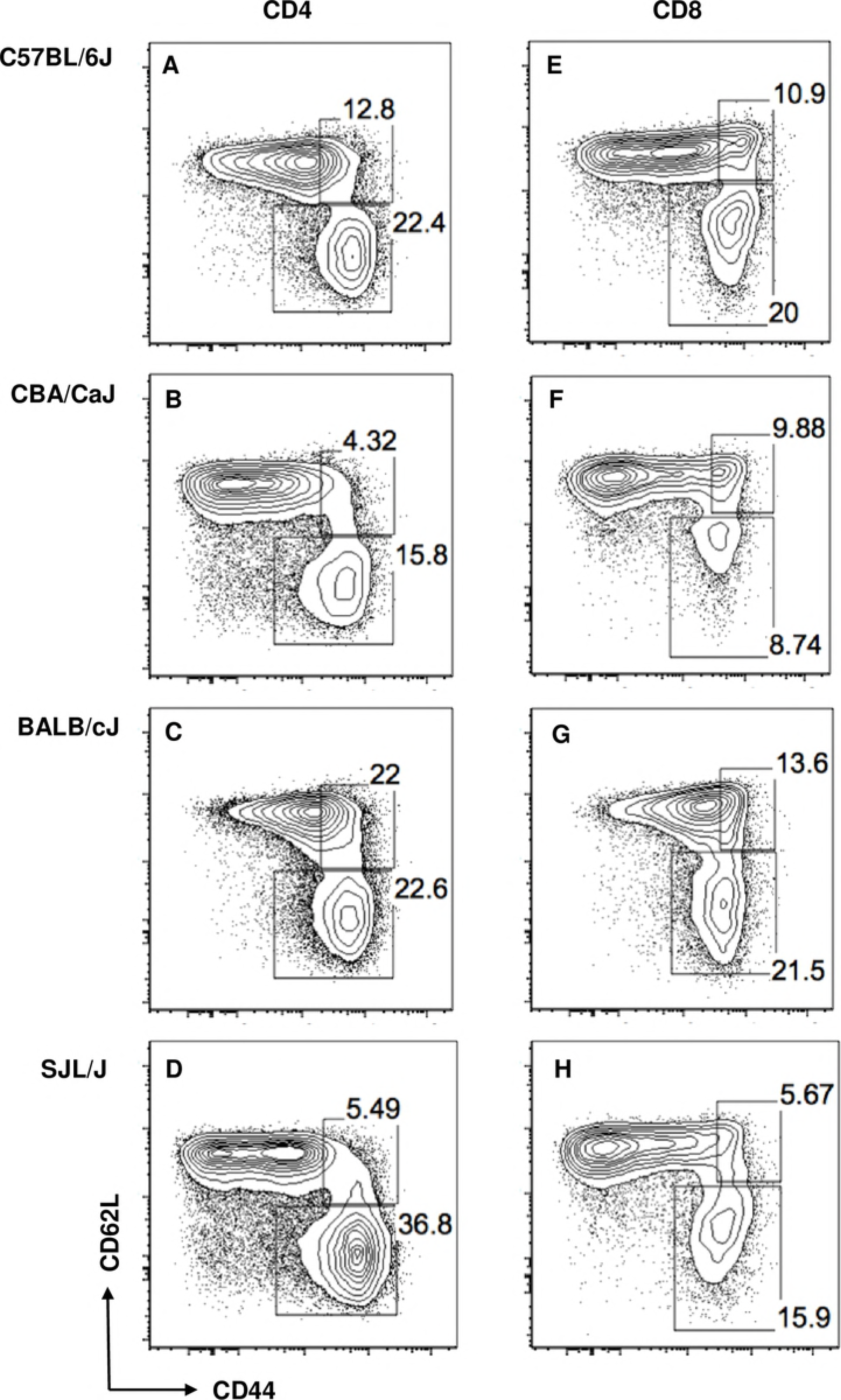
Frequencies of EMCD4 and EMCD8 in the spleen from donor partner in mixed bone marrow chimera of B6.SJL-CBA/CaJ pair (A, B) and BALB/cJ-SJL/J pair (C, D) Each dot represents data from one individual mouse (n > 9 per group) and individual mice data are connected by lines. The p-values are as indicated in the panels.

### CD4 gene expression in strains that differ in CD4 effector memory phenotype

Since bone marrow chimera data suggested cell-autonomous regulation of memory phenotype, we hypothesized that intrinsic gene expression differences in naive CD4 T cells between the mouse strains could contribute to differences in EMCD4 frequencies that are observed across the strains. We tested this using gene expression data (microarray) of splenic CD4 T cells from multiple mouse strains available from the immunological genome project (33). Immgen gene expression data comes from sorted CD4+ CD62L+ spleen cells (‘non-EM’ CD4 T cells), which could potentially include CD44-high CMCD4 population in addition to genuine naïve CD4 T cells. Hence, we selected 2 mouse strains (CBA/CaJ and SJL/J) that showed maximum differences in EMCD4 frequencies and counts, without showing differences in CMCD4 levels (Fig S4 A to E, p-values shown in supplementary table S5). CMCD4 frequencies were very low in these 2 strains (Figure 3B and 3D), making comparison of CD4+ CD62L+ gene expression between these strains interpretable. A comparison of gene expression profile of CD4+ CD62L+ve splenocytes (obtained from GEO accession: GSE60337) of CBA/CaJ and SJL/J identified 290 genes that were significantly different (p values < 0.05 after multiple correction) (supplementary table S6). Gene ontology enrichment analysis identified significantly enriched GO processes associated with the 290-gene list (supplementary table S6). As expected, MHC molecule related transcripts were differentially expressed and came up as significantly enriched, reassuring that the analysis we performed could actually pick up biologically relevant differences (table S6). In addition, there was significant enrichment for lipid biosynthesis (11 genes, p value: 1.78E-05, adjusted p value: 0.004) and lipid metabolism (16 genes, p value: 1.13E-04, adjusted p value: 0.02) (supplementary file S6), implying that these pathways could be of biological relevance to regulation of CD4 T cell memory phenotype. When data from multiple strains were pooled together, there was a positive correlation between EMCD4 and EMCD8 frequencies (Fig S4F), similar to what is observed for humans (Fig 2B), suggesting similar mechanisms in regulation of CD4 and CD8 effector memory or dependence of one subset on the other.

## Discussion

There are a number of possible explanations for the observation of a substantial extent of variation in the relative frequencies of memory-phenotype B cells, CD4 T cells and CD8 T cells. Such variations in biological parameters are commonly interpreted as errors in technical or experimental factors, although in real populations, biological variance may well have meaningful interpretative value (31). At the population level, we have used descriptive characterization of the human peripheral blood leucocyte phenotype to examine the possible significance of population level variation in these leucocyte subsets.

By comparison of intra-individual variance of memory and effector subsets of T and B cells, we find that immune subsets that respond to day-to-day fluctuations, such as the plasmablasts and CD4 TEMRA cells, show similar intra-individual and inter-individual variance in the population, while B cell memory, CD4 memory and CD8 memory cell frequencies are very much stable over a one-year period within individuals. This suggests that memory subsets are not determined by short-term environmental fluctuations, unlike CD4 TEMRA and plasmablasts. Within-individual variation can thus possibly be an explanatory factor in the population-level variation for effector T cells and plasmablast subsets, but not for memory T and B cell subsets. A recent study (45) that quantified technical and biological variation in human immune phenotype also found high intra-individual variation in CD4 TEMRA cells in comparison to other immune lineages. Our data show that CD8 TEMRA cells did not behave similar to CD4 TEMRA cells in that they do not show considerable intra-individual fluctuation. This could be related to indications that CD8 TEMRA cells may be functionally different from CD4 TEMRA cells, despite being phenotypically similar (46).

Our findings of a lack of correlation between B cell memory and CD4 memory also argue against the possibility of cumulative exposures determining both the memory subsets, although it remains possible that cumulative exposures regulate one but not the other subset. The positive correlation we find between CD4 memory and CD8 memory suggest that T cell memory levels could be regulated by similar mechanisms, distinct from how memory B cell levels are regulated.

We attempted to quantify the degree of similarity in immune phenotype between siblings to see if there was any indication that these memory phenotypes are heritable. Although our sibling study was not as powerful as previously published twin studies (1,2,4,14) for understanding heritability, our sibling data strongly support and extend the interpretations from our serial bleed data. Thus, siblings did not show concordance in effector T and B subsets, suggesting that variation in those subsets is environmentally driven. On the other hand, siblings were far similar to each other in memory subset frequencies than non-siblings were, consistent with a genetic component regulating memory lymphocyte frequencies. However, it must be acknowledged that any similarity between siblings could be attributed to early life influences, since it is well recognized that both intra-uterine (47,48) and neonatal (49) stress impacts immune system development, and a recent study (13) has shown a prominent role for co-habitation in determining immune phenotypes. Our findings are consistent with two previous studies (1,14) which report high degree of heritability for majority of immune subsets. Another study (2), too, reported high degree of heritability for CD4 memory subsets, although it found non-heritable factors to be important for most other immune phenotypes. Since our interpretations are predominantly based on associations, we complemented the study with experimental data from mice.

Although laboratory animals show restricted genetic diversity compared to human populations, we exploited the fact that independent inbred strains are genetically distinct and are grown in a homogeneous environment in a controlled animal facility. Hence, it is worth examining if differences in leucocyte subset levels between strains are genetically driven. We immunophenotyped some common strains of mice and chose two pairs showing substantial differences, SJL/J and BALB/cJ for further experiments on CD4 memory and C57BL/6J and CBA/CaJ for CD8 and CD4 T cell memory phenotype. It is notable that not only did splenic CD4 and CD8 memory frequencies show differences, but that total numbers of these cells per organ also showed the same differences, indicating that subset frequencies are reasonable surrogates for the pool size of these subsets. We also have noted that animals maintained in our facility had relatively high frequencies of EM phenotype T cells even at steady state. Even though mice were harbored in a specific-pathogen-free facility, it remains possible that there might be variations in microbial antigenic burden or gut microbiome composition between laboratories, although these factors are difficult to quantify. Long years of reproductive isolation because of inbreeding could also have contributed to these differences.

Our mixed bone marrow chimera experiments allowed us to examine whether the CD4 and/or CD8 memory levels were genetically determined and whether these genetic influences were T cell-intrinsic in nature. Our data suggest that donor genotype-specific cell-intrinsic factors strongly influence both CD4 and CD8 memory T cell pool size.

Using gene expression data of mouse splenic CD4+ve CD62L+ve T cells available in the public domain (33) we attempted to characterize genetic differences that could explain differences in CD4 memory phenotype. There are a number of limitations in our approach. Firstly, we do not evaluate gene expression differences in the same mice (or even mice from the same small animal facility) that we experimentally find phenotypic differences for, and differences in phenotype that we observe could be influenced by additional facility-specific environmental factors as well.

Secondly, the array based gene expression data available in the public domain that we used (33) contain only 2 replicates for each strain (except for C57BL/6J), limiting statistical power and increasing the chances of picking up false positive differences. Thirdly, the “naïve” population as defined by Immgen is based on CD4+ve CD62L+ve gate, and does not exclude CD44+ cells, potentially including central memory cells into the population. Our analysis attempts to overcome this limitation by comparing those strains which differ only in EMCD4, and not CMCD4 subsets. In spite of these caveats, our analysis is still useful as a preliminary exploration that can be extended in further experiments using multiple replicates and strains. Remarkably, lipid/fatty acid metabolism pathways that came up as significant in as limited an analysis as this, was recently suggested to play crucial roles in T cell activation (50). This exploratory analysis is thus likely to provide further clues to direct future mechanistic studies.

Thus, our data suggest genetic and cell-intrinsic factors as major determinants of the memory pool size of T cells as well as, probably, B cells. However, in a natural ecosystem where animals are exposed to substantial and continual antigenic exposures, the balance between cell-intrinsic genetic effects on memory and the effects of environmental factors is likely to be nuanced and quantitative. Further studies using varying antigenic burdens and experimental immunizations will be necessary to address these issues. Mechanistically, it will be interesting to explore whether genetic cell-intrinsic factors affect memory generation during immune response or memory attrition after the peak of immune response. Our data provide interesting insights and directions for future work in understanding the genesis and consequences of the regulation of niche size of lymphocyte memory.

## Acknowledgements

We acknowledge Department of Biotechnology, Government of India and Department of Science and Technology, Government of India for funding the study.

## Funding

The study was supported in part by grants from the Department of Biotechnology (to A.G. # BT/PR12849/MED/15/35/2009; to V.B. # BT/PR14420/Med/29/213/2010; to S.R. # BT/PR-14592/BRB/10/858/2010; to S B. BT/MB/01/THSTI-ChBC/2009; and to U.C.M.N. # BT/PR14723/MED/15/44/2010), and from the Department of Science and Technology, Government of India (to V.B. # SR/SO/HS-0005/2011 and #EMR/2015/001074; to S.R. # SB/SO/HS/210/2013). The National Institute of Immunology and the Translational Health Science and Technology Institute are supported by the Department of Biotechnology, Government of India.

## Conflict of interest

The authors declare no conflict of interest.

## List of abbreviations

EM: : Effector memory
TEMRA: : T, effector memory, RA positive
CD: : Cluster of differentiation

